# Fatty links between multisystem proteinopathy and Small VCP-Interacting Protein

**DOI:** 10.1101/2023.03.31.531359

**Authors:** Firyal Ramzan, Fatima Abrar, Ashish Kumar, Lucia Meng Qi Liao, Zurie E. Campbell, Rachel V. Gray, Oluwadurotimi Akanni, Colm Guyn, Dale D. O. Martin

## Abstract

Multisystem proteinopathy (MSP) is a rare dominantly-inherited disorder that includes a cluster of diseases, including frontotemporal dementia, inclusion body myopathy, and Paget’s disease of bone. MSP is caused by mutations in the gene encoding Valosin-Containing Protein (VCP). Patients with the same mutation, even within the same family, can present with a different combination of any or all of the above diseases, along with amyotrophic lateral sclerosis (ALS). The pleiotropic effects may be linked to the greater than 50 VCP cofactors that direct VCP’s many roles in the cell. Small VCP-Interacting Protein (SVIP) is a small protein that directs VCP to autophagosomes and lysosomes. We found that SVIP directs VCP localization to autophagosomes in an acylation-dependent manner. We demonstrate that SVIP is myristoylated at glycine 2 and palmitoylated at cysteines 4 and 7. Acylation of SVIP is required to mediate cell death in the presence of the MSP-associated VCP variant (R155H-VCP), whereas blocking SVIP myristoylation prevents cytotoxicity. Therefore, SVIP acylation may present a novel target in MSP.

## INTRODUCTION

Point mutations in the gene encoding for Valosin Containing Protein (VCP) have been causally and correlatively linked to numerous autosomal dominant neurodegenerative diseases. These include VCP diseases and multisystem proteinopathy (MSP), in which patients may present with one or any combination of inclusion body myopathy, Paget’s disease of bone, fronto-temporal dementia, amyotrophic lateral sclerosis (ALS) (Scarian et al., 2022), Parkinson’s disease (Kirby et al., 2021), and others (Al-Obeidi et al., 2018).

VCP is a homohexameric AAA-ATPase involved in many cellular functions, including endoplasmic reticulum-associated degradation (ERAD) and autophagy, that are regulated by its cofactors (Hänzelmann & Schindelin, 2017). We directed our studies toward its cofactors to understand VCP’s role in disease. One of these cofactors is Small VCP-interacting protein (SVIP) is a VCP cofactor that regulates VCP’s function in activation of autophagy and negatively regulate ERAD (Ballar et al., 2007; Jia et al., 2019; Johnson et al., 2021; Wang et al., 2011). SVIP is a small protein that binds the N-terminal domain of VCP (Hänzelmann & Schindelin, 2011; Nagahama et al., 2003), and can redirect VCP localization to autophagosomes (Wang et al., 2011). Recent studies in *Drosophila melanogaster* showed that reducing SVIP disrupts muscle lysosomal function, and causes muscular and neuromuscular degeneration, motoneuronal degeneration, motor dysfunction, and reduced lifespan (Johnson et al., 2021). These studies further characterized an SVIP mutation identified in patients with sporadic frontotemporal dementia and confirmed its pathogenicity in *Drosophila melanogaster* (Johnson et al., 2021).

Previous studies have linked SVIP localization and function to N-myristoylation (Ballar et al., 2007; Nagahama et al., 2003), the non-reversible addition of the 14-carbon saturated fatty acid myristate to N-terminal glycines by N-myristoyl transferase (NMT) enzymes 1 and 2 (NMT1 and NMT2) (Martin et al., 2011; Meinnel et al., 2020). N-myristoylation, herein referred to as myristoylation, can occur either co-translationally, after the removal of the initiator methionine on the nascent polypeptide, or post-translationally, following the exposure of an N-terminal glycine of the C-terminal product of proteolyzed proteins. The myristate moiety promotes weak membrane binding and often requires a second membrane binding site for stable membrane attachment. Therefore, myristoylated proteins typically undergo additional forms of lipidation at downstream sites. In the case of SVIP, there are two cysteines immediately downstream of the glycine that are predicted to undergo S-acylation. Similar to myristoylation, S-acylation involves the addition of saturated fatty acids, but via a thioester bond on cysteine residues, making it labile and reversible. The 16-carbon fatty acid palmitate is typically used and, therefore, this modification is more commonly referred to as S-palmitoylation or simply palmitoylation.

Although there is no known consensus sequence for palmitoylation, N-and C-terminal cysteines are often palmitoylated, especially when adjacent to a myristoylated site. In contrast, myristoylation typically occurs on N-terminal glycines found within a GXXXT/S/C consensus motif, where X can be almost any amino acid. Prolines and large bulky amino acids are not usually tolerated in these positions. Therefore, SVIP does not have an ideal myristoylation consensus sequence (MGLCFPCPG). Regardless, SVIP is still predicted to be myristoylated. Previous reports that the N-terminus is required for SVIP localization suggested SVIP is indeed fatty acylated (Ballar et al., 2007; Nagahama et al., 2003). However, no studies have confirmed either myristoylation or palmitoylation of SVIP and, importantly, their role in regulating SVIP function as a VCP cofactor. We hypothesized that SVIP is palmitoylated at cysteines 4 and 7, and myristoylated at glycine 2.

In this study, we confirm SVIP is both myristoylated and palmitoylated. Moreover, we find that localization of both SVIP and VCP is modulated by SVIP acylation. Further, SVIP has an acylation-dependent toxic interaction with R155H-VCP, a disease-associated variant of VCP.

## METHODS

### Cell culture and transfection

HEK293T cells were grown at 37˚C in 5% CO_2_ in Dulbecco’s Modified Eagle Medium (DMEM; Wisent # 319- 005-CL, supplemented with 10% FBS, 0.1% Penicillin-streptomycin, 0.1% L-Glutamine, 0.1% sodium pyruvate). Cells were seeded at 300,000 cells/well for biochemical experiments or 225,000 cells/well for microscopy experiments in 6-well dishes. The next day, cells were transfected with 5 µg of plasmid DNA using calcium phosphate precipitation as previously described (Martin et al., 2019). Cells were transfected for 2 h, followed by a media change, and were processed for analysis 18 h later.

### Mouse-derived Primary Hippocampal Neurons

Primary hippocampal neurons were derived from mouse (FVBN/J strain) embryos at embryonic day 17 (E17). Embryonic hippocampi were dissociated using papain and seeded at 180,000 cells/well for microscopy experiments on 0.1 mg/mL poly-L-lysine coated wells or glass coverslips (#1.5, VWR) in 6-well plates. They were plated in plating media (Neurobasal supplemented with 2.5% FBS, 2.5% Horse serum, pen-strep, L-Glutamine, and B27). The next day, the media was changed to neuronal media (Neurobasal supplemented with pen-strep, LL-glutamine, and B27). At 14 days in vitro, cells were transfected with lipofectamine 2000 (Invitrogen) and fixed the following day using 4% paraformaldehyde in 1x PBS.

### Plasmids

SVIP is predicted to be myristoylated at Glycine 2 the prediction programs TermiNator (https://bioweb.i2bc.paris-saclay.fr/terminator3/test.php) and SVMyr (https://busca.biocomp.unibo.it/lipipred/3a05fb48-b5c8-40f5-93e0-71427220d8bf/showresult/). SVIP is predicted to be palmitoylated at Cysteines 4 and 7 by CSS Palm 3.0 (http://csspalm.biocuckoo.org/index.php; (Ren et al., 2008)). All SVIP-mCherry plasmids were generated using gBlock or gene strands (IDT or Eurofins, respectively). Briefly, the open reading frame of SVIP was transferred from the plasmid into FEW-mCherry viral vector (kind gift from Dr. Gareth Thomas, Temple University) using the XhoI and NotI restrictions enzyme sites, resulting in a C-terminal mCherry tag. The resulting constructs are referred to as WT-SVIP-mCherry in the following text. G2A, C4S, C7S, C4,7S, and G2A-C4,7S-SVIP-mCherry refer to the SVIP site mutants in the FEW-mCherry vector. Additionally, a gBlock was used to induce 3 site mutations (R22E, A26V, R32E) in the VCP-interacting motif (VIM) region in WT-SVIP-mCherry (VIMm-SVIP-mCherry) (See Figure 1a for a summary of the site mutations).

**Figure 1.**
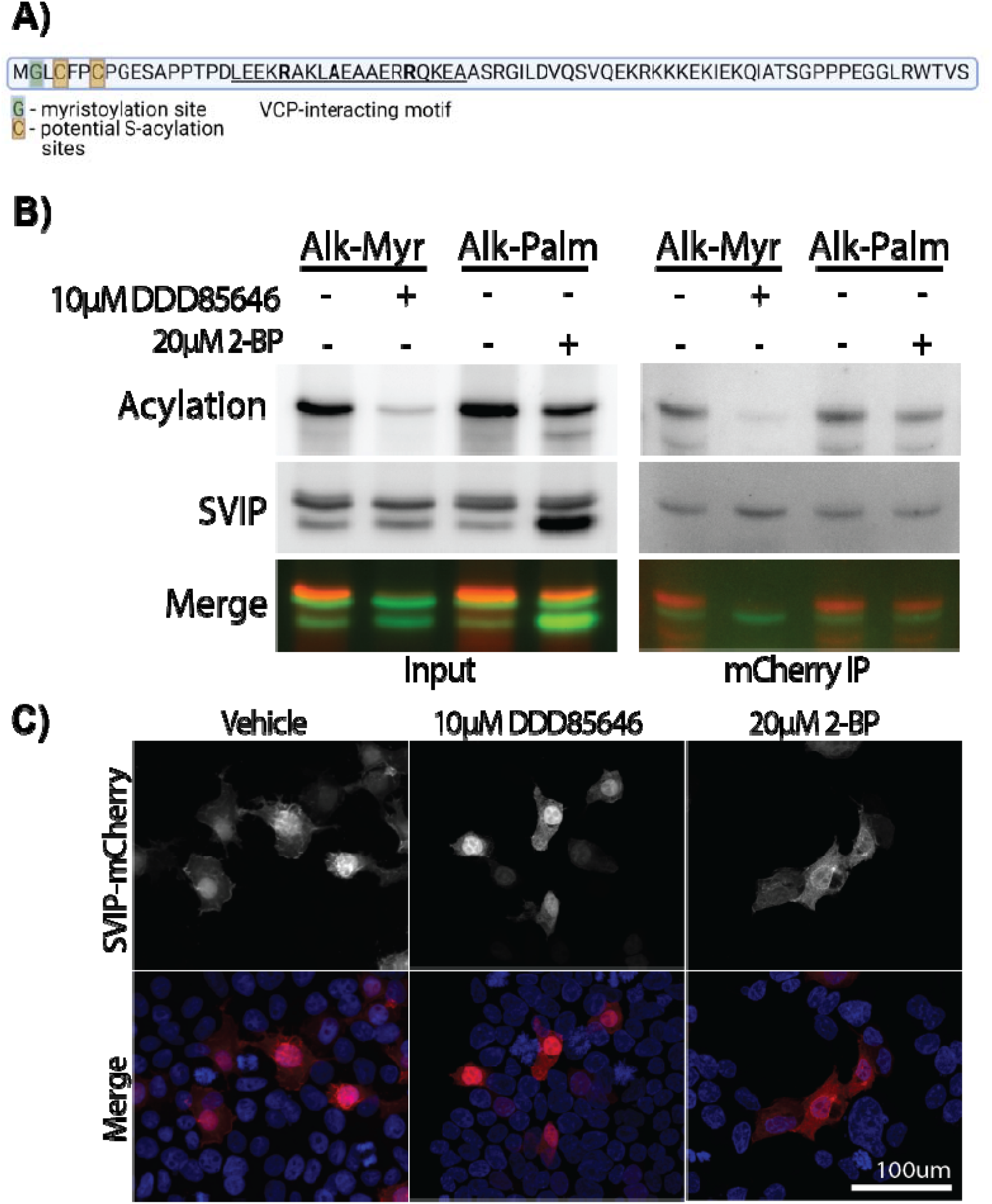
SVIP is myristoylated and palmitoylated. A) Amino Acid sequence of SVIP. SVIP is a small 77 amino acid long protein that is predicted to undergo myristoylation at Glycine 2 (G; Green; TermiNator and SVMyr) and palmitoylation at Cysteines 4 and 7 (C; orange; CSS-Palm 3.0). The VCP binding motif (VIM) is underlined and essential amino acids are bolded. B) SVIP-mCherry incorporation of alkyne-myristate and alkyne-palmitate was detected by click chemistry and is sensitive to their respective myristoylation (DDD85646) and palmitoylation (2- BP) inhibitors in HEK293T cells. C) Myristoylation or palmitoylation inhibition alters SVIP-mCherry localization in HEK293T cells.

The open reading frames of WT-VCP-eGFPN1, R155H-VCP-eGFPN1, and A232E-VCP-eGFPN1 were transferred from the eGFP plasmid into FEW-GFP viral vector using standard molecular cloning techniques with the NotI and SalI restriction enzyme sites, resulting in a C-terminal GFP tag. The resulting plasmids were used for all experiments and are referred to as WT-VCP-GFP and R155H-VCP or R155H-VCP-GFP in the following text.

### Fatty acid labelling and Click Chemistry

Cell labelling and click chemistry were performed as described previously (Liao et al., 2021; Yap et al., 2010). Briefly, cells were deprived of lipids for 45 min – 1 h in DMEM supplemented with delipidated FBS. Then, 15 min prior to labelling, cells were treated with the indicated inhibitors [1 µM DDD85646 dissolved in DMSO or 20 µM 2-bromopalmitate (2BP) dissolved in ethanol] where indicated. Next, alkyne-tagged fatty acid analogs (alkyne-myristate (13-tetradecylnoic acid; 13-TDYA; Click Chemistry Tools 1164) or alkyne-palmitate (15-hexadecynoic acid; 15-HDYA; Click Chemistry Tools 1165) were saponifed in potassium hydroxide. Cells were then labelled with the saponified alkyne fatty acid analogs for 3 hr, after which cells were lysed in modified RIPA buffer (50 mM HEPES pH 7.4, 150 mM NaCl, 0.5% sodium deoxycholate, 2 mM MgCl_2_, 0.1% SDS, 1% Igepal CA-630). Rabbit anti-mcherry (Cell Signaling) and goat anti-GFP (Eusera) were used to immunoprecipitate (IP) SVIP-mCherry and VCP-GFP, respectively. IPs and lysates (input) then underwent a click reaction with tris-carboxyethylphosphine (TCEP), copper sulphate, S-(benzyltriazolylmethyl) amine (TBTA), and azide-647.

### Acyl-biotin exchange (ABE)

The ABE was performed as previously described (Hurst et al., 2017; Martin et al., 2019). Briefly, cells were lysed and sonicated in lysis buffer (50mM HEPES, 2% SDS, 1mM EDTA) with 50 mM N-ethylmaleimide (NEM). Lysates were incubated in a 50°C water bath for 2 h to block free thiols with NEM. NEM was then removed during a 2 h incubation at room temperature, rotating in the presence of 2,3-dimethyl-1,2-butadiene, added to a final concentration of 4%, followed by chloroform precipitation of the butadiene (final concentration of 8%) and phase separation of excess NEM. The top phase was collected and incubated with 0.7 M hydroxylamine and 4 mM HPDP-Biotin (Soltec) for 1 hr rotating at room temperature to cleave thioester bonds. After an overnight 80% acetone precipitation at -20˚C, protein pellets were washed with 80% acetone, and resuspended in lysis buffer without NEM. Following protein quantification, 500 µg – 1 mg of protein was incubated in 150mM NaCl dilution buffer (50 mM HEPES, 1% TX-100, 1 mM EDTA, 1 mM EGTA) with High-Capacity Neutravidin Agarose beads (Thermo #29202) at 4°C for 4 h. Following two washes with 0.5 M NaCl dilution buffer and 1 wash with dilution buffer without NaCl, the protein was eluted from beads with elution buffer (0.2% SDS, 250 mM NaCl, 1% β-mercaptoethanol) over a 10 m incubation at 37˚C, then denatured at 95˚C for 5 m in 5X sample-loading buffer with 5% β-mercaptoethanol, and stored at -20°C.

### Western blotting

10% Bis-Tris acrylamide gels were run in XT MOPS buffer (Biorad #1610788) and transferred onto nitrocellulose using the Transblot Turbo (Biorad). Rat anti-mCherry (1:1000, Invitrogen), and rabbit anti-GFP (1:10,000, Eusera) were used to detect SVIP-mCherry and VCP-GFP, respectively. Fluorescent secondaries anti-mouse Starbright 700 (Biorad) and anti-rabbit Starbright 520 (Biorad) were used. Tubulin-rhodamine (Biorad) was used to detect Tubulin as the loading control.

### Fluorescence microscopy

HEK293T cells were seeded at 225,000 cells/well onto 0.01 mg/mL poly-D-lysine coated coverslips (#1.5, VWR) in 6-well plates. After an 18 h transfection (described above), cells were washed once with 1X phosphate-buffered saline (PBS), then fixed in 4% paraformaldehyde for 20 m at room temperature. After 2 PBS washes, cells were incubated in 1µg/mL DAPI (Sigma D9542) in PBS for 20 m at room temperature, followed by 3 PBS washes. The coverslips were mounted onto microscope slides (VWR 48311-601) using ProLong Gold antifade mountant (Invitrogen P36934), and sealed with nail polish. Images were acquired using the Nikon AXR laser scanning confocal microscope at 20X magnification. Using Nikon AXR software, z-stack images at 20X were taken, which were first denoised, followed by deconvolution, and the resulting maximum intensity projections are reported below.

Hippocampal neurons seeded at 180,000 cells/well onto 0.1 mg/mL poly-L-lysine coated coverslips (#1.5, VWR) in 6-well plates. After an 18 h transfection, cells were fixed in 4% paraformaldehyde for 20 min as described above. Images were acquired using the Nikon AXR laser scanning confocal microscope at 60X magnification. Z-stacks images were taken, which were denoised, followed by deconvolution, and the resulting maximum intensity projections are reported below.

For live cell imaging, HEK293T cells were plated at approximately 225,000 cells/well on poly-D-Lysine coated 1.5 coverglass in 35 mm live-cell culture dishes (MatTek). The next day, cells were transfected with the indicated constructs. After approximately 2 hours, media was replaced and cells were incubated with LysoTracker Deep Red. In order to capture dying cells, cells expressing R155H-VCP-GFP and WT-SVIP-mCherry were imaged overnight at 60X magnification and maintained at 37°C at 5% CO_2_. All other conditions were imaged the next day under the same conditions.

## RESULTS

### SVIP is both myristoylated and palmitoylated

SVIP has 3 predicted acylation sites, G2, C4, C7 (Figure 1A). Using click chemistry, we confirmed that SVIP-mCherry is both myristoylated and palmitoylated (Figure 1B). The acylation signal was also prevented by pretreatment with the acylation inhibitors 15 minutes before alkyne-fatty acid labelling. Specifically, treatment with the myristoylation inhibitor DDD85646 reduced SVIP myrisotylation with the the alkyne-myristate label, while treatment with the palmitoylation inhibitor 2-bromopalmitate (2-BP) reduced SVIP palmitoylation with the alkyne-palmitate label (Figure 1B).

When expressed alone, SVIP-mCherry appears to localize to membranes throughout the cell (Figure 1C). However, upon treatment with DDD85646, SVIP-mCherry becomes diffusely expressed throughout the cell. Like other short acylated peptides appended to a fluorescent protein (Martin et al., 2008, 2012; McCabe & Berthiaume, 1999, 2001), blocking myristoylation results in SVIP-mCherry localizing similar to mCherry alone with more nuclear localization. Blocking SVIP-mCherry palmitoylation with 2-BP only partially altered SVIP-mCherry localization (Figure 1C). This is likely because myristoylation is required for palmitoylation of SVIP-mCherry, and blocking myristoylation with DDD85646 completely blocks all SVIP-mCherry acylation (both palmitoylation and myristoylation) while blocking palmitoylation with 2-BP only blocks palmitoylation, allowing SVIP-mCherry to partially localize to membranes. Thus, SVIP-mCherry, follows the standard pattern of fatty acylation at the N-terminus (Martin et al., 2008, 2012; McCabe & Berthiaume, 1999, 2001).

### SVIP is myristoylated at Glycine 2, and palmitoylated at Cysteines 4 and 7

To confirm myristoylation is required for palmitoylation and to verify the sites of acylation, we introduced an alanine mutation at G2 (G2A) and serine mutations at C4 and C7 (C4S and C7S) and detected myristoylation and palmitoylation (Figure 2A-B). Serine is more sterically similar to cysteine than alanine. Using alkyne-myristate labelling and click chemistry, we confirmed that SVIP-mCherry myristoylation requires the essential glycine at position 2 (G2A and G2A-C4,7S). While the C7S substitution did not affect myristoylation, the C4S and C4,C7S mutations consistently had a lower myristoylation signal compared to WT (Figure 2A). This suggests that the cysteine in position 4 is likely required for recognition by NMT. To ensure that the substitution to serine was not the cause, we also generated a cysteine to alanine mutation that also reduced myristoylation (data not shown) indicating that cysteine at position 4 is required for myristoylation of SVIP-mCherry.

**Figure 2.**
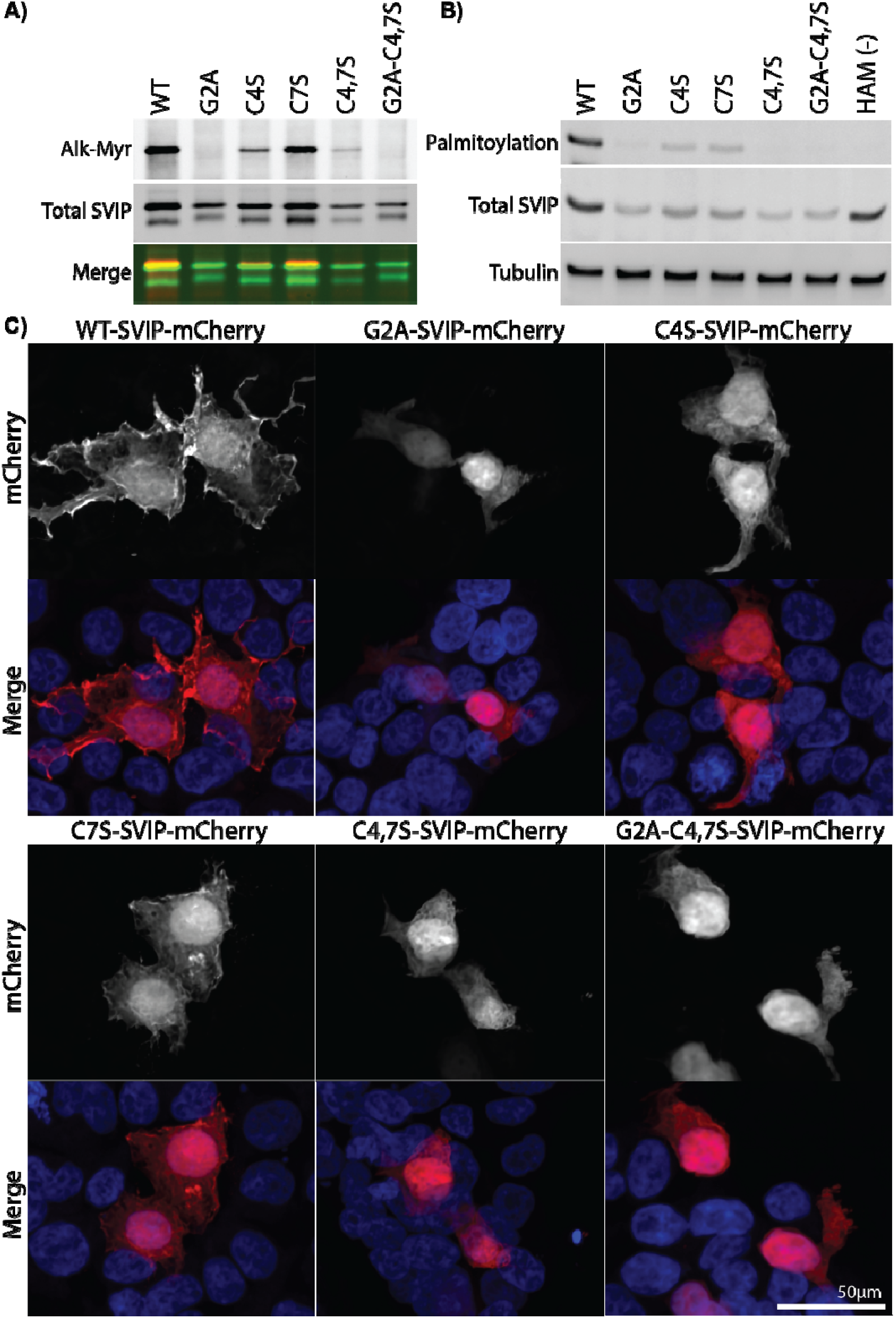
SVIP is myristoylated at Glycine 2 and palmitoylated at Cysteines 4 and 7. Point mutations at the indicated positions were made to block myristoylation (G2) and palmitoylation (C4,7S) in SVIP-mCherry. HEK293T cells expressing these mutations were used to detect A) Myristoylation by click chemistry using alkyne-myristate and B) palmitoylation by ABE. C)SVIP-mCherry localization was acquired by fixed cell confocal microscopy.

To assess palmitoylation more directly, an acyl-biotin exchange (ABE) assay was performed. As predicted, point mutations at C4 and C7 reduced SVIP palmitoylation, but not completely, whereas the double mutation (C4,7S) completely blocked palmitoylation, indicating SVIP-mCherry is palmitoylated at C4 and C7. Palmitoylation was nearly completely inhibited when myristoylation was blocked in G2A (Figure 2B), confirming that myristoylation is required for downstream palmitoylation. The faint signal that remained is likely due to partial palmitoylation at the most distal C7 site. This confirms that myristoylation is required for SVIP palmitoylation.

We next wanted to determine if SVIP’s acylation regulates its localization in HEK293T cells. WT-SVIP is localized throughout the cell and on membrane ruffles, whereas non-acylated SVIP (G2A) is more diffuse within the cell and nuclear (Figure 2C). Substitution at the C4 site had a similar effect, likely due to the reduced myristoylation and palmitoylation. C7S-SVIP had an intermediate effect with some membrane-specific localization, but not as prominent as WT-SVIP. The combined mutants, C4,7S and G2A-C4,7S, both demonstrate a general diffuse pattern of localization, likely due to the presence of the G2A and/or C4S mutations (Figure 2C), which block myristoylation and palmitoylation.

### SVIP-mCherry directs VCP-GFP localization and is SVIP acylation-dependent

To assess how acylation may regulate SVIP co-localization with VCP, SVIP-mCherry and VCP-GFP were co-expressed in HEK293T cells (Figure 3A). Alone, WT-VCP-GFP was diffusely expressed throughout the cell. When expressed with WT-SVIP-mCherry, both VCP-GFP and SVIP-mCherry localize to vesicles reminiscent of autophagosomes (Supplemental Video 1), as previously described (Wang et al., 2011). Strikingly, the formation of these autophagosomes was dependent on SVIP-mCherry acylation. No vesicles formed in the presence of G2A-SVIP-mCherry (Supplemental Video 2). However, regardless of the acylation mutant, VCP-GFP and SVIP-mCherry localization overlapped, suggesting the proteins still interact.

**Figure 3.**
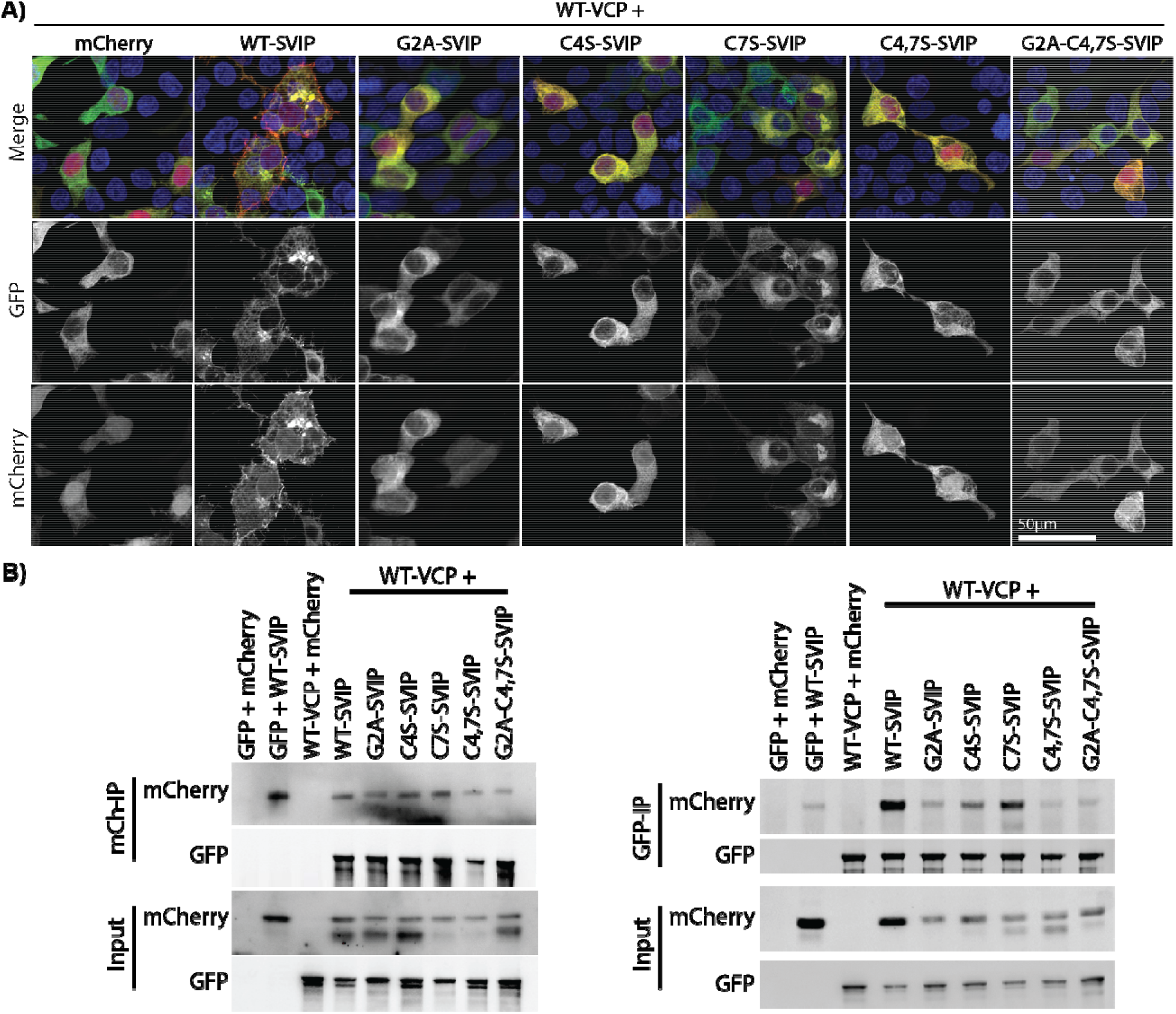
SVIP acylation is required to direct VCP localization. A) HEK293T cells co-expressing WT-VCP-GFP and SVIP-mCherry acylation mutants were imaged by confocal microscopy. B) Co-immunoprecipitation with anti-mCherry (*left*) and anti-GFP (*right*) from cell lysates of HEK293T cells expressing the combinations of WT-VCP-GFP with SVIP-mCherry acylation mutants.

To confirm the interaction, we performed co-immunoprecipitation studies using WT-VCP-GFP and the acylation mutants of SVIP-mCherry (Figure 3B). When immunoprecipitating SVIP-mCherry, the acylation status did not appear to affect the co-immunoprecipitation of VCP-GFP (Figure 3B, left panel). In contrast, when VCP-GFP was immunoprecipitated, SVIP-mCherry mutants that had reduced or no myristoylation (G2A, C4,7S, and G2A-C4,7S), and therefore little to no palmitoylation, were reduced to background levels (Figure 3B, right panel). This suggests VCP is a primary interactor of SVIP and the interaction is independent of acylation. However, because VCP has many interactors, non-acylated SVIP may not have the opportunity to interact with VCP and, therefore, less co-immunoprecipitates with VCP.

### SVIP-mCherry and R155H-VCP-GFP have an acylation-dependent toxic interaction

Next, we sought to determine if the acylation-dependent effect of SVIP and VCP was maintained in VCP disease. The R155H-VCP variant accounts for almost 50% of VCP disease cases, while the A232E variant is linked to more aggressive VCP phenotypes (Tresse et al., 2010). Surprisingly, when R155H-VCP-GFP was co-expressed with WT-SVIP-mCherry, we detected rapid cell deterioration, ultimately leading to cell death soon after expression of both proteins (Supplemental Video 3). These findings were replicated biochemically (Figure 4A). Specifically, protein levels of both R155H-or A232E-VCP-GFP and WT-SVIP-mCherry were substantially reduced when co-expressed. Furthermore, this rapid cell death was resistant to caspase inhibition (Figure 4B), suggesting caspase-independent cell death. To determine if this toxicity was acylation dependent, we co-expressed R155H-VCP-GFP with acylation mutants of SVIP-mCherry. Notably, blocking SVIP acylation (G2A, C4S, C4,7S, and G2A-C4,7S) rescued the toxic effect as measured biochemically (Figure 4C) and by microscopy (Figure 4D), with the exception of C7S-SVIP-mCherry, which appeared to have an intermediate effect.

**Figure 4.**
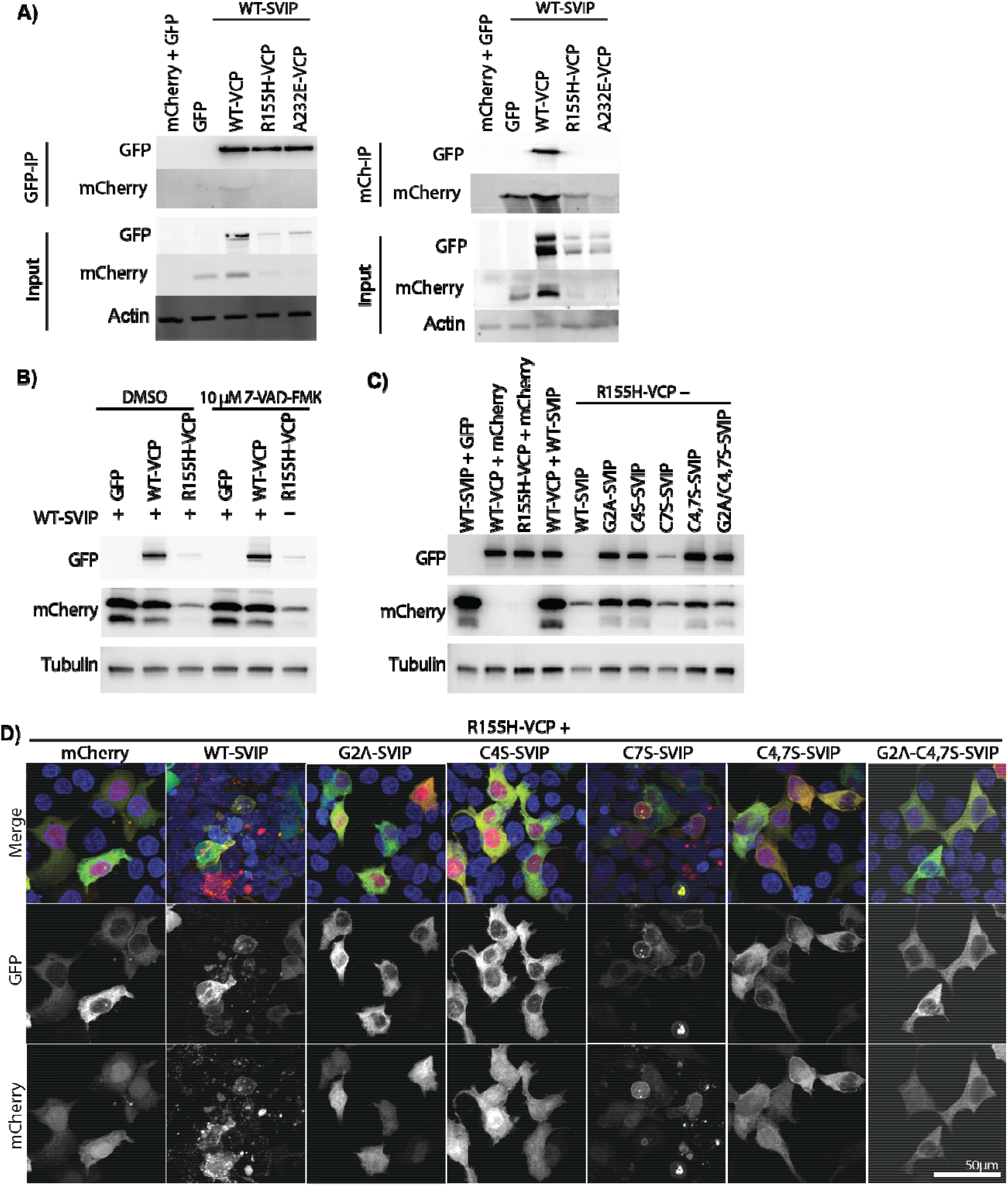
The toxic interaction of SVIP and R155H-VCP is rescued by blocking SVIP acylation. A) Coexpression of SVIP-mCherry with mutant VCP-GFP (R155H or A232E) was toxic to HEK293T cells and led to the degradation of the two proteins. B) Toxicity was not rescued by caspase inhibition with the general caspase inhibitor Z-VAD-FMK. The toxic effects were reversed when acylation was blocked and visualized C) biochemically or D) by confocal microscopy.

### Treatment with acylation inhibitors rescues toxicity

Because the C4S substitution in SVIP also decreased myristoylation (Figure 2A), it was difficult to determine if the protective effect of blocking acylation was mediated by palmitoylation or myristoylation. This was compounded by the fact that myristoylation is required for downstream palmitoylation at C4 and C7 (Figure 2B). To determine if palmitoylation was necessary, cells co-expressing R155H-VCP-GFP and WT-SVIP-mCherry were treated with 10 µM DDD85646 or 20 µM 2-BP to inhibit myristoylation or palmitoylation, respectively. Treatment with the myristoylation inhibitor (DDD85646) completely prevented the toxicity (Figure 5). In contrast, treatment with the palmitoylation inhibitor (2-BP) appeared to have an intermediate effect on the toxic interaction such that, upon fixation, cells were found rounding up (Figure 5A). These findings were recapitulated biochemically (Figure 5B). Specifically, protein levels of both R155H-VCP-GFP and WT-SVIP-mCherry were high when myristoylation was inhibited (G2A-SVIP, DDD85646 treatment), and substantially reduced when treated with vehicle (DMSO) or the palmitoylation inhibitor (2-BP) (Figure 5B). This suggests that the toxicity observed upon coexpression of R155H-VCP-GFP and WT-SVIP-mCherry is largely dependent on myristoylation.

**Figure 5.**
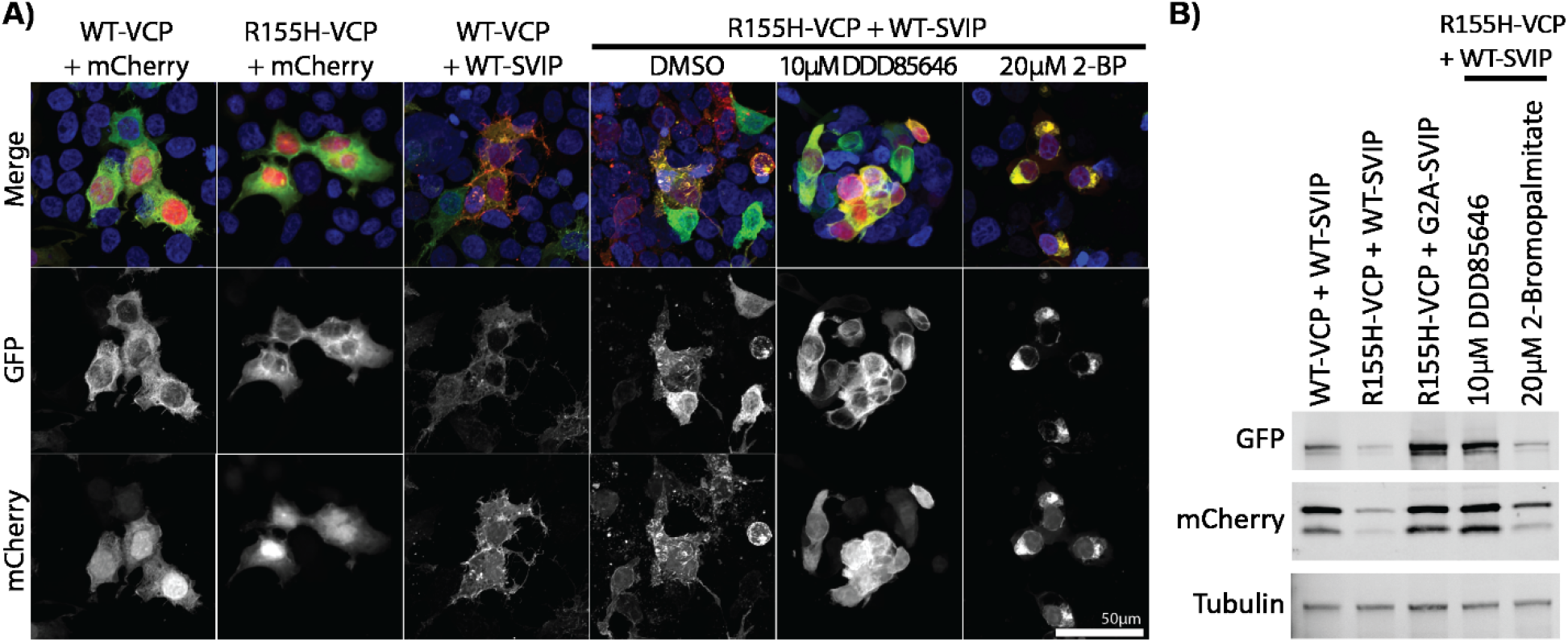
Global inhibition of palmitoylation or myristoylation rescues toxicity. HEK293T cells co-transfected with R155H-VCP-GFP and WT-SVIP-mCherry were treated with vehicle (DMSO), 10 µM DDD85646, and 20 µM 2- BP overnight for 18hr. A) Cells treated with inhibitors were imaged by fixed cell confocal microscopy. B) Western blot results depict the rescue in protein expression levels when treated with DDD85646 and moderate rescue with 2-BP treatment.

### Blocking the SVIP VCP-interaction motif (VIM) rescues toxicity

To determine if the toxic effect was dependent on the interaction of VCP and SVIP, 3 point mutations were made to the VIM region (R22E, A26V, R32E^18^LEEKRAKLAEAAERRQKE^35^, Figure 1A) in SVIP-mCherry (VIMm-SVIP-mCherry). As previously published (Ballar et al., 2007), this caused SVIP-mCherry to migrate slightly higher by SDS-PAGE and was sufficient to prevent WT-VCP-GFP from co-immunoprecipitating with SVIP-mCherry (Figure 6A). However, VIMm-SVIP-mCherry had a reduced but still prominent interaction with R155H-VCP-GFP that was only detected in the GFP immunoprecipitate, but not the mCherry (Figure 6A). This reduction was sufficient to rescue toxicity (Figure 6A and B), indicated by microscopy and biochemically. By microscopy, VIMm-SVIP-mCherry still localized to intracellular membranes (Figure 6B, top row), reminiscent of WT-SVIP-mCherry (Figures 1-5) and did not overlap with R155H-VCP-GFP, which was dispersed throughout the cell (Figure 6B, bottom row), similar to when it is expressed on its own (Figure 6B, second row).

**Figure 6.**
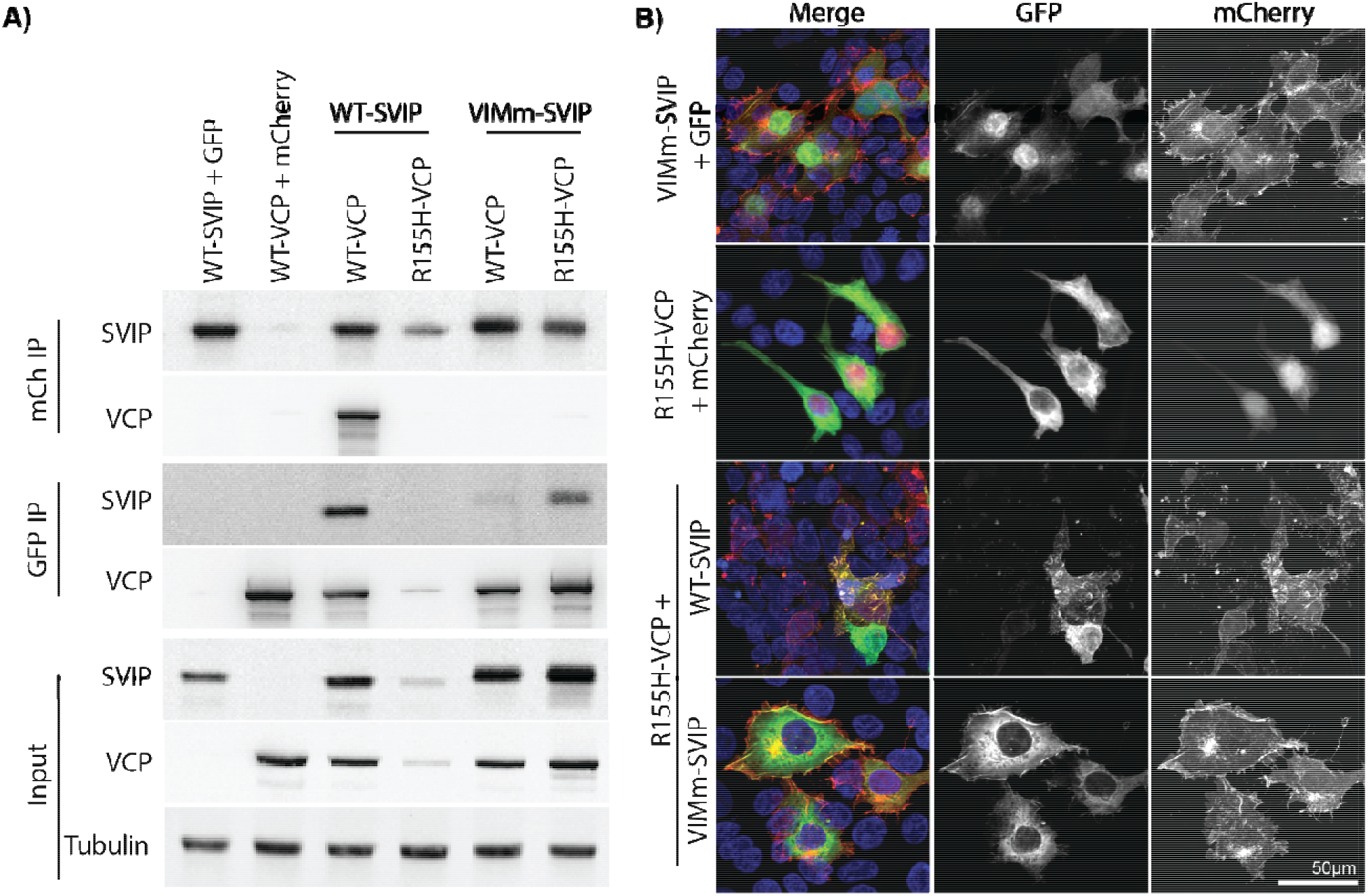
Blocking SVIP’s interaction with VCP with the VIM mutation rescues toxicity. SVIP-mCherry with multiple site mutations in the VCP-interacting motif (VIMm-SVIP-mCherry) was generated and co-expressed with R155H-VCP-GFP in HEK293T cells. A) Co-immunoprecipitation results depict that the VIMm-SVIP-mCherry does not interact with WT-VCP, but does with R155H-VCP and rescues R155H-VCP protein expression. B) Fixed cells were imaged using the Nikon AXR confocal microscope at 20X magnification.

### Lysosome localization & toxicity is dependent on SVIP acylation

Using live-cell microscopy (Figure 7, Supplemental Videos 4-7), we see that acylation of WT-SVIP is required to localize SVIP-mCherry and WT-VCP-GFP to Lysotracker stained lysosomes (Figure 7A, Supplemental Video 4), as previously published. When SVIP acylation is blocked (G2A-SVIP-mCherry), the two proteins no longer localize to the lysosome and appear diffuse within the cytoplasm, suggesting SVIP is required to target the two proteins to the lysosome (Figure 7A, Supplemental Videos 5 and 6). However, when WT-SVIP-mCherry is co-expressed with R155H-VCP-GFP (Figure 7B, Supplemental Video 7), the cells die soon after VCP expression is detected (approximately 60-100 mins). Initially, WT-SVIP-mCherry can be seen localizing to the lysosomes in cells not expressing R155H-VCP-GFP. Once GFP is detected, the cells quickly begin dying. No apparent mutant VCP is detected at the lysosome with WT-SVIP-mCherry. Again, this toxicity was reversed after blocking SVIP acylation and localization to the lysosome. This suggests that SVIP must localize to the lysosome to mediate toxicity in the presence of mutant VCP.

**Figure 7.**
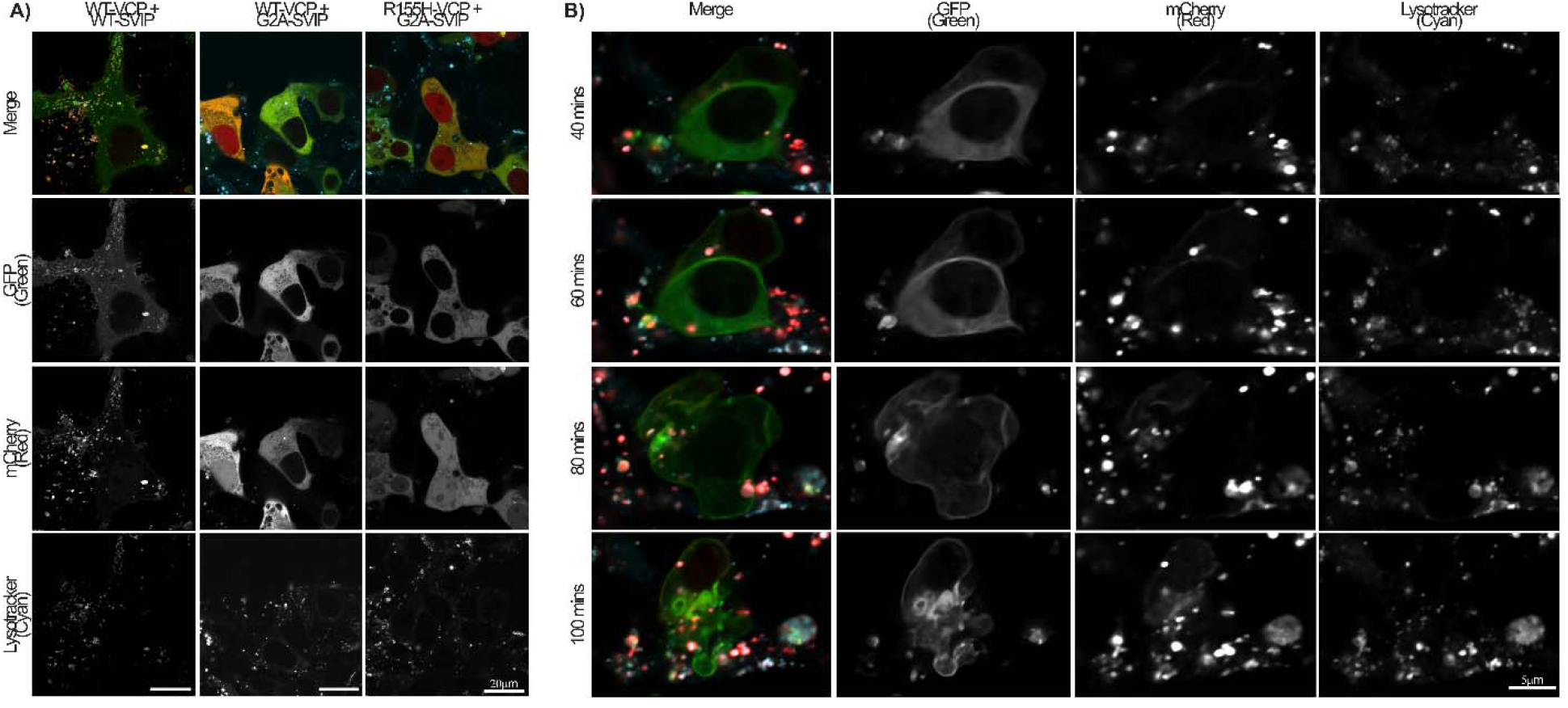
Lysosome targeting and toxicity is acylation-dependent. HEK293T expressing WT or G2A-SVIP-mCherry with WT or R155H-VCP-GFP were incubated with Lysotracker Deep Red (Cyan) soon after transfection and imaged A) the next day or B) overnight using the Nikon AXR confocal microscope at 60X magnification.

### VCP and SVIP localization is replicated in neurons

Mouse-derived primary hippocampal neurons were transfected with SVIP-mCherry and WT-VCP-GFP (Figure 8). As seen in HEK293T cells, WT-SVIP-mCherry was found along membranes, particularly along the axons and dendrites, while G2A-SVIP-mCherry was nuclear and dispersed diffusely throughout the cell body, axons, and dendrites. Again, coexpression of WT-VCP-GFP with WT-SVIP-mCherry results in autophagosome-like vesicles found in the cell body and throughout the neurites (Figure 8, panel 3 inset), which are no longer detected with coexpression of G2A-SVIP-mCherry and WT-VCP-GFP. Blocking SVIP-mCherry acylation resulted in a general diffuse expression pattern of both proteins.

**Figure 8.**
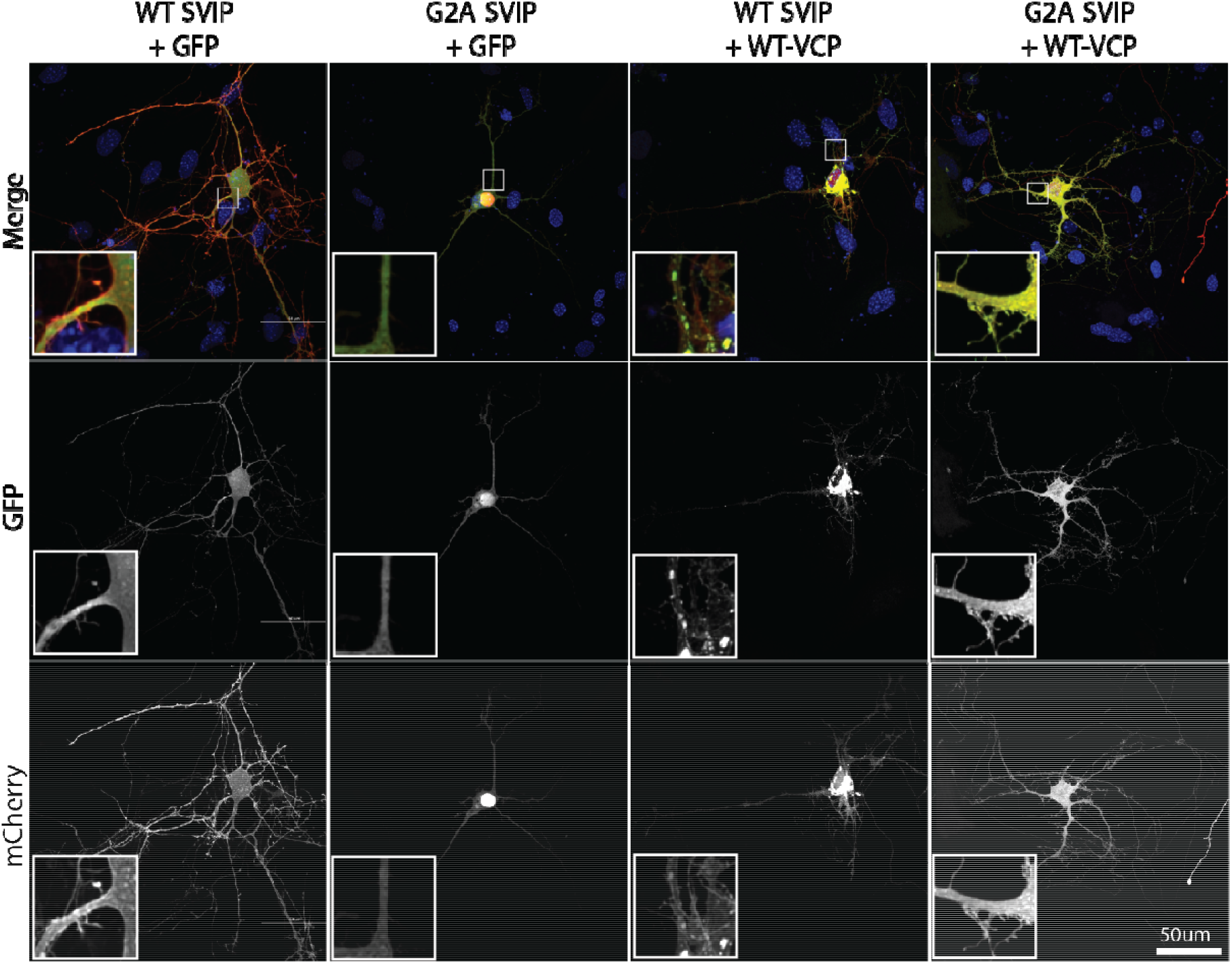
SVIP-mCherry and VCP-GFP expression in hippocampal neurons. Primary hippocampal neurons derived from FVB mice were transfected with WT-or G2A-SVIP-mCherry and GFP or WT-VCP-GFP. Neurons were imaged by fixed cell confocal microscopy at 60X. Insets depict a 50% zoomed-in view of neurites at the base of the cell body.

## DISCUSSION

Previous studies have indicated that the N-terminus of SVIP is required for SVIP localization and affects VCP localization (Wang et al., 2011). Here we confirm that human SVIP is myristoylated at G2, which is required for the downstream palmitoylation at sites C4 and, to a lesser degree, C7 (Figure 2). In turn, SVIP acylation is required for SVIP localization and can regulate VCP localization when co-expressed. Together, as previously indicated (Wang et al., 2011), SVIP-mCherry and VCP-GFP localize to autophagosome-like vesicles and is dependent on SVIP-mCherry acylation (Figure 3, Supplemental Videos 1 and 2). Furthermore, the combination of SVIP with disease variants of VCP is lethal to the cell through a caspase-independent form of cell death that is SVIP acylation-dependent.

We found that SVIP’s cellular localization is differentially regulated by myristoylation and palmitoylation. Specifically, blocking myristoylation via the G2A mutation resulted in the relocalization of SVIP from membranes to diffused throughout the cell (Figure 2), including the nucleus. This effect was replicated with a myristoylation inhibitor (Figure 1B). Nuclear localization was possibly an effect of the fluorescent tag in combination with blocking myristoylation and palmitoylation. As seen in other proteins that are dually acylated at the N-terminus, myristoylation at G2 is required for downstream palmitoylation (at C4 and C7). When short acylated peptides are appended to fluorescent proteins and acylation is inhibited, the proteins localize diffusely in the cytoplasm, as well as the nucleus (Martin et al., 2008, 2012; McCabe & Berthiaume, 1999, 2001). Thus, the G2A mutation blocks nearly all acylation of SVIP-mCherry, leading the protein to localize like mCherry alone. Alternatively, SVIP may contain a nuclear localization signal that directs non-acylated SVIP to the nucleus.

Interestingly, the C4S substitution has similar effects on SVIP-mCherry localization as the G2A mutation (Figure 2). This is likely due to the decreased myristoylation detected by click chemistry in the C4S mutation (Figure 2B). Substitution of C4 to alanine did not restore myristoylation (data not shown). As previously mentioned, SVIP does not have a canonical myristoylation consensus motif, which may make it more sensitive to substitutions at the N-terminus. As expected, blocking palmitoylation at C7 did not alter myristoylation (Figure 2A) and had only an intermediate effect on localization (Figure 2C).

Prior evidence suggests that SVIP directs VCP localization to the plasma membrane and lysosomes (Wang et al., 2011). Here we confirm that VCP-GFP localization is dependent on SVIP-mCherry acylation (Figure 3A). Together, the two proteins appear to induce the formation of autophagosomes in an SVIP acylation-dependent mechanism. Again, there is an intermediate effect when only palmitoylation at C7 is blocked (C7S), which leads to SVIP-mCherry and VCP-GFP becoming concentrated along the edge of the nucleus rather than a general diffuse pattern as with other acylation mutants (Figure 3A), in contrast to the partial membrane localization of C7S-SVIP-mCherry alone (Figure 2C).

However, the reduced signal of SVIP acylation mutants upon co-immunoprecipitation of VCP suggests reduced interaction of both G2A-and C4S-SVIP-mCherry with VCP-GFP (Figure 3B). It is likely that the interaction between SVIP and VCP may be reliant on SVIP localization and, thus, acylation status. SVIP is the only VCP cofactor known to assemble with each VCP subunit of the VCP hexamer (6 SVIP:6 VCP monomers) (Hänzelmann & Schindelin, 2011, 2017). Therefore, immunoprecipitation of any SVIP is likely to co-immunoprecipitate VCP, but SVIP co-immunoprecipitation with VCP may not be detected due to competition with many other cofactors.

In an effort to confirm our results with VCP mutants linked to multisystem proteinopathy, we found that R155H-VCP-GFP has a rapid toxic interaction with WT-SVIP-mCherry, wherein expression of both proteins results in cell death soon after expression (Supplemental Figure 3). This toxicity is prevented when SVIP-mCherry acylation is blocked via both site mutations or with biochemical inhibitors, suggesting that SVIP acylation impacts its interaction with the disease-causing mutant of VCP. There is an intermediate attenuation of toxicity with C7S and 2-BP treatment, indicating blocking myristoylation may be required for complete prevention of toxicity. Specifically, there may be reduced interaction of R155H-VCP with G2A-and C4S-SVIP, resulting in the prevention of toxicity. As mentioned previously, there is some indication in the co-immunoprecipitation results that the acylation mutants of SVIP may have reduced interaction with VCP. This may carry over with R155H-VCP as well and rescue the toxic interaction. Indeed, when the VCP-interaction motif mutant of SVIP (VIMm-SVIP) was co-expressed with R155H-VCP, it failed to co-immunoprecipitate with both wild-type and R155H-VCP (Figure 6a). VIMm-SVIP also rescued the toxic phenotype when co-expressed with R155H-VCP (Figure 6). This further supports the hypothesis that the interaction between R155H-VCP and SVIP may be modulated through SVIP myristoylation, and, perhaps, through SVIP palmitoylation at C4.

The toxic phenotype observed between R155H-VCP-GFP and WT-SVIP-mCherry in our studies contrasts previous findings (Johnson et al., 2021). In *Drosophila melanogaster*, SVIP knockout results in significant muscular, mitochondrial, and neuronal degeneration while restoring muscular SVIP expression rescues many of these degenerative effects, including neuronal degeneration (Johnson et al., 2021). This may be explained by a combination of factors. First, our system uses overexpression of mCherry tagged SVIP and VCP-GFP. However, the toxicity was confined to the disease variant R155H and not observed with WT-VCP-GFP. Cell death was also observed when SVIP-mCherry was co-expressed with R155H-VCP-GFP in multiple cell types (HEK293T, HeLa, and primary hippocampal neurons – not shown). Finally, toxicity was highly specific to SVIP-mCherry acylation status, which likely differs in *Drosophila melanogaster*. In flies, SVIP has an additional cysteine at position 8 as well as a more canonical myristoylation site (MGAC<u>LS</u>CCGQ…), in comparison to mammalian SVIP, which only has cysteines at positions 4 and 7, and a non-canonical myristoylation site (MGLC<u>FP</u>CPGE…). Specifically, F5, P6, and P8 in human SVIP are not ideal amino acids for myristoylation. This suggests that human SVIP is not fully myristoylated, and therefore palmitoylated, and may exist in multiple forms based on its acylation status (i.e. there may be pools of acylated and non-acylated SVIP). Thus, *Drosophila melanogaster* SVIP is likely a better substrate for myristoylation and more efficiently palmitoylated, due to the third cysteine, than mammalian SVIP, affecting its localization and function differently than in mammals. Additionally, while VCP is highly conserved from yeast to humans (White & Lauring, 2007), SVIP does not appear to be as well conserved between species (35% sequence identity between fly and human SVIP in contrast to approximately 85% for VCP (The UniProt Consortium et al., 2023)). This difference in conservation may be reflective of diverse functions in different species. It may suggest that as humans developed disease variants of VCP over time, SVIP acylation became less desirable. As such, it will be critical to study the role of SVIP acylation in multiple species and living systems to develop a comprehensive understanding of SVIP’s complex role in disease. That said, based on the requirement of SVIP for tubular lysosomes in *Drosophila* and those in our study, SVIP may be a membrane-curvature sensing and inducing protein that is dependent on acylation. Many membrane curvature sensing and inducing proteins are fatty acylated (Hatzakis et al., 2009; Larsen et al., 2015; Martin et al., 2014). Therefore, the potentially more efficient myristoylation and additional palmitoylation of SVIP in *Drosophila* may make SVIP a better curvature inducing protein to facilitate the formation of tubular lysosomes.

The mechanism behind this toxic interaction needs to be explored in future experiments. One potential mechanism is alterations in autophagy. It is important to note that autophagosome-like vesicles are produced when the wild-type forms of SVIP and VCP are co-expressed in multiple tested cell types (primary hippocampal neurons (Figure 8 panel 3), HEK293T (Figure 2A), and HeLa (not shown)). Data from previous literature indicates that these vesicles are indeed autophagosomes (Wang et al., 2011). Since the interaction between WT-VCP and WT-SVIP results in autophagosomes and SVIP regulates VCP’s functions in autophagy, this process may become aberrantly regulated with R155H-VCP and WT-SVIP, rapidly deteriorating the cells expressing both proteins. Other possible mechanisms of toxicity include ER-associated degradation and apoptosis, both of which VCP is typically involved in (Meyer & Weihl, 2014). However, the resistance to caspase inhibitors suggests a caspase-independent form of cell death.

The regulation of VCP and SVIP interaction by acylation will be a critical avenue to explore. The results of this study demonstrate the importance of both SVIP myristoylation and palmitoylation in modulating its regulation of VCP localization and the resulting toxicity with the most common disease-causing variant of VCP.

## Acknowledgements

We would like to thank Cure VCP Disease Inc. for hosting the Cure VCP Disease Research forum and introducing us to an international research group and would like to extend our gratitude to members of this forum for their input and support. This work was supported by a Natural Sciences and Engineering Research Council (NSERC) Discovery Grant (RGPIN-2019-04617)to DDOM and a Mitacs Accelerate Grant in collaboration with Circumvent Pharmaceuticals Inc. FR is currently supported by Canadian Institutes of Health Research (CIHR 472640) Postdoctoral Award, and LMQL is funded through an ALS Canada Trainee Award.

## References

Al-Obeidi, E., Al-Tahan, S., Surampalli, A., Goyal, N., Wang, A. K., Hermann, A., Omizo, M., Smith, C., Mozaffar, T., & Kimonis, V. (2018). Genotype-phenotype study in patients with valosin-containing protein mutations associated with multisystem proteinopathy. Clinical Genetics, 93(1), 119–125. https://doi.org/10.1111/cge.13095

Ballar, P., Zhong, Y., Nagahama, M., Tagaya, M., Shen, Y., & Fang, S. (2007). Identification of SVIP as an endogenous inhibitor of endoplasmic reticulum-associated degradation. The Journal of Biological Chemistry, 282(47), 33908–33914. https://doi.org/10.1074/jbc.M704446200

Hänzelmann, P., & Schindelin, H. (2011). The Structural and Functional Basis of the p97/Valosin-containing Protein (VCP)-interacting Motif (VIM). Journal of Biological Chemistry, 286(44), 38679–38690. https://doi.org/10.1074/jbc.M111.274506

Hänzelmann, P., & Schindelin, H. (2017). The Interplay of Cofactor Interactions and Post-translational Modifications in the Regulation of the AAA+ ATPase p97. Frontiers in Molecular Biosciences, 4(APR), 1–22. https://doi.org/10.3389/fmolb.2017.00021

Hatzakis, N. S., Bhatia, V. K., Larsen, J., Madsen, K. L., Bolinger, P.-Y., Kunding, A. H., Castillo, J., Gether, U., Hedegård, P., & Stamou, D. (2009). How curved membranes recruit amphipathic helices and protein anchoring motifs. Nature Chemical Biology, 5(11), 835–841. https://doi.org/10.1038/nchembio.213

Hurst, C. H., Turnbull, D., Plain, F., Fuller, W., & Hemsley, P. A. (2017). Maleimide scavenging enhances determination of protein S-palmitoylation state in acyl-exchange methods.BioTechniques, 62(2), 69–75. https://doi.org/10.2144/000114516

Jia, D., Wang, Y. Y., Wang, P., Huang, Y., Liang, D. Y., Wang, D., Cheng, C., Zhang, C., Guo, L., Liang, P., Wang, Y., Jia, Y., & Li, C. (2019). SVIP alleviates CCl4-induced liver fibrosis via activating autophagy and protecting hepatocytes. Cell Death & Disease, 10(2), 71. https://doi.org/10.1038/s41419-019-1311-0

Johnson, A. E., Orr, B. O., Fetter, R. D., Moughamian, A. J., Primeaux, L. A., Geier, E. G., Yokoyama, J. S., Miller, B. L., & Davis, G. W. (2021). SVIP is a molecular determinant of lysosomal dynamic stability, neurodegeneration and lifespan. Nature Communications, 12(1), 513. https://doi.org/10.1038/s41467-020-20796-8

Kirby, A. E., Kimonis, V., & Kompoliti, K. (2021). Ataxia and Parkinsonism in a Woman With a VCP Variant and Long-Normal Repeats in the SCA2 Allele. Neurology Genetics, 7(4), e595. https://doi.org/10.1212/NXG.0000000000000595

Larsen, J. B., Jensen, M. B., Bhatia, V. K., Pedersen, S. L., Bjørnholm, T., Iversen, L., Uline, M., Szleifer, I., Jensen, K. J., Hatzakis, N. S., & Stamou, D. (2015). Membrane curvature enables N-Ras lipid anchor sorting to liquid-ordered membrane phases. Nature Chemical Biology, 11(3), 192–194. https://doi.org/10.1038/nchembio.1733

Liao, L. M. Q., Gray, R. A. V., & Martin, D. D. O. (2021). Optimized Incorporation of Alkynyl Fatty Acid Analogs for the Detection of Fatty Acylated Proteins using Click Chemistry.Journal of Visualized Experiments, 2021(170), 1–17. https://doi.org/10.3791/62107

Martin, D. D. O., Ahpin, C. Y., Heit, R. J., Perinpanayagam, M. A., Yap, M. C., Veldhoen, R. A., Goping, I. S., & Berthiaume, L. G. (2012). Tandem reporter assay for myristoylated proteins post-translationally (TRAMPP) identifies novel substrates for post-translational myristoylation: PKC_∊_, a case study. The FASEB Journal, 26(1), 13–28. https://doi.org/10.1096/fj.11-182360

Martin, D. D. O., Beauchamp, E., & Berthiaume, L. G. (2011). Post-translational myristoylation: Fat matters in cellular life and death. Biochimie, 93(1), 18–31. https://doi.org/10.1016/j.biochi.2010.10.018

Martin, D. D. O., Heit, R. J., Yap, M. C., Davidson, M. W., Hayden, M. R., & Berthiaume, L. G. (2014). Identification of a post-translationally myristoylated autophagy-inducing domain released by caspase cleavage of Huntingtin. Human Molecular Genetics, 23(12), 3166–3179. https://doi.org/10.1093/hmg/ddu027

Martin, D. D. O., Kanuparthi, P. S., Holland, S. M., Sanders, S. S., Jeong, H.-K., Einarson, M. B., Jacobson, M. A., & Thomas, G. M. (2019). Identification of Novel Inhibitors of DLK Palmitoylation and Signaling by High Content Screening. Scientific Reports, 9(1), 3632. https://doi.org/10.1038/s41598-019-39968-8

Martin, D. D. O., Vilas, G. L., Prescher, J. A., Rajaiah, G., Falck, J. R., Bertozzi, C. R., & Berthiaume, L. G. (2008). Rapid detection, discovery, and identification of post-translationally myristoylated proteins during apoptosis using a bio-orthogonal azidomyristate analog. The FASEB Journal, 22(3), 797–806. https://doi.org/10.1096/fj.07-9198com

McCabe, J. B., & Berthiaume, L. G. (1999). Functional Roles for Fatty Acylated Amino-terminal Domains in Subcellular Localization. Molecular Biology of the Cell, 10(11), 3771–3786. https://doi.org/10.1091/mbc.10.11.3771

McCabe, J. B., & Berthiaume, L. G. (2001). N-Terminal Protein Acylation Confers Localization to Cholesterol, Sphingolipid-enriched Membranes But Not to Lipid Rafts/Caveolae. Molecular Biology of the Cell, 12(11), 3601–3617. https://doi.org/10.1091/mbc.12.11.3601

Meinnel, T., Dian, C., & Giglione, C. (2020). Myristoylation, an Ancient Protein Modification Mirroring Eukaryogenesis and Evolution. Trends in Biochemical Sciences, 45(7), 619–632. https://doi.org/10.1016/j.tibs.2020.03.007

Meyer, H., & Weihl, C. C. (2014). The VCP/p97 system at a glance: Connecting cellular function to disease pathogenesis. Journal of Cell Science, 127(18), 3877–3883. https://doi.org/10.1242/jcs.093831

Nagahama, M., Suzuki, M., Hamada, Y., Hatsuzawa, K., Tani, K., Yamamoto, A., & Tagaya, M. (2003). SVIP Is a Novel VCP/p97-interacting Protein Whose Expression Causes Cell Vacuolation.Molecular Biology of the Cell, 14(1), 262–273. https://doi.org/10.1091/mbc.02-07-0115

Ren, J., Wen, L., Gao, X., Jin, C., Xue, Y., & Yao, X. (2008). CSS-Palm 2.0: An updated software for palmitoylation sites prediction. Protein Engineering Design and Selection, 21(11), 639–644. https://doi.org/10.1093/protein/gzn039

Scarian, E., Fiamingo, G., Diamanti, L., Palmieri, I., Gagliardi, S., & Pansarasa, O. (2022). The Role of VCP Mutations in the Spectrum of Amyotrophic Lateral Sclerosis—Frontotemporal Dementia.Frontiers in Neurology, 13(February), 1–15. https://doi.org/10.3389/fneur.2022.841394

The UniProt Consortium, Bateman, A., Martin, M.-J., Orchard, S., Magrane, M., Ahmad, S., Alpi, E., Bowler-Barnett, E. H., Britto, R., Bye-A-Jee, H., Cukura, A., Denny, P., Dogan, T., Ebenezer, T., Fan, J., Garmiri, P., da Costa Gonzales, L. J., Hatton-Ellis, E., Hussein, A., … Zhang, J. (2023). UniProt: The Universal Protein Knowledgebase in 2023. Nucleic Acids Research, 51(D1), D523–D531. https://doi.org/10.1093/nar/gkac1052

Tresse, E., Salomons, F. A., Vesa, J., Bott, L. C., Yao, T., Dantuma, N. P., Taylor, J. P., Tresse, E., Salomons, F. A., Vesa, J., Bott, L. C., Yao, T., Dantuma, N. P., Taylor, J. P., Tresse, E., Salomons, F. A., Vesa, J., Bott, L. C., Kimonis, V., … Dantuma, N. P. (2010). VCP/p97 is essential for maturation of ubiquitin-containing autophagosomes and this function is impaired by mutations that cause IBMPFD.Autophagy, 6(2), 217–227. https://doi.org/10.4161/auto.6.2.11014

Wang, Y., Ballar, P., Zhong, Y., Zhang, X., Liu, C., Zhang, Y.-J., Monteiro, M. J., Li, J., & Fang, S. (2011). SVIP Induces Localization of p97/VCP to the Plasma and Lysosomal Membranes and Regulates Autophagy.PLoS ONE, 6(8), e24478. https://doi.org/10.1371/journal.pone.0024478

White, S. R., & Lauring, B. (2007). AAA+ ATPases: Achieving Diversity of Function with Conserved Machinery. Traffic, 8(12), 1657–1667. https://doi.org/10.1111/j.1600-0854.2007.00642.x

Yap, M. C., Kostiuk, M. A., Martin, D. D. O., Perinpanayagam, M. A., Hak, P. G., Siddam, A., Majjigapu, J. R., Rajaiah, G., Keller, B. O., Prescher, J. A., Wu, P., Bertozzi, C. R., Falck, J. R., & Berthiaume, L. G. (2010). Rapid and selective detection of fatty acylated proteins using ω-alkynyl-fatty acids and click chemistry.Journal of Lipid Research, 51(6), 1566–1580. https://doi.org/10.1194/jlr.D002790

